# Erp and Rev adhesins of the Lyme disease spirochete’s ubiquitous cp32 prophages assist the bacterium during vertebrate infection

**DOI:** 10.1101/2022.12.01.518731

**Authors:** Brian Stevenson, Catherine A. Brissette

**Author notes:** Addresses for correspondence: Brian Stevenson, Department of Microbiology, Immunology, and Molecular Genetics, University of Kentucky College of Medicine, Lexington, Kentucky, 40536-0298, Phone: 1+859-257-9358, Catherine A. Brissette, Department of Biomedical Sciences, School of Medicine and Health Sciences, University of North Dakota, Grand Forks, North Dakota, 58203-9061, Phone: 1+701-777-6412.

## Abstract

Almost all spirochetes in the genus *Borrelia* (sensu lato) naturally contain multiple variants of closely related prophages. In the Lyme disease borreliae, these prophages are maintained as circular episomes that are called cp32s (<u>c</u>ircular <u>p</u>lasmid <u>32</u>kb). The cp32s of Lyme agents are particularly unique in that they encode two distinct families of lipoproteins, Erp and Rev, that are expressed on the bacteria’s outer surface during infection of vertebrate hosts. All identified functions of those outer surface proteins involve interactions between the spirochetes and host molecules: Erp proteins bind plasmin(ogen), laminin, glycosaminoglycans, and/or components of complement, and Rev proteins bind fibronectin. Thus, cp32 prophages provide their bacterial hosts with surface proteins that can enhance infection processes, thereby facilitating their own survival. Horizontal transfer via bacteriophage particles increases spread of beneficial alleles and creates diversity among Erp and Rev proteins.

## Introduction

Members of the spirochete genus *Borrelia* (sensu lato) persist through infectious cycles between blood-feeding arthropods and vertebrates, and include agents of infections that plague humans, domestic animals, and wildlife (1, 2). These include Lyme disease, caused by *Borrelia burgdorferi* sensu lato, and tick-borne relapsing fever, caused by *B. hermsii, B. turicatae*, and others. Although a proposal has been raised to divide the genus into *Borrelia* for relapsing fever agents and *Borreliella* for Lyme disease agents, that idea is controversial (3-8). For the purpose of this review, we use the unified genus name *Borrelia*, in part because of the shared property among Lyme disease and relapsing fever spirochetes of near-universal natural colonization with multiple, closely-related cp32 prophages (9-13) (Figs. 1 and 2). Of particular significance to this review, the cp32s of both Lyme disease and relapsing fever borreliae carry *mlp* genes, while *erp* and *rev* genes appear to be restricted to the Lyme disease borreliae (11). The significance of those genes will be discussed further below. For simplicity, we also use the name *Borrelia burgdorferi* to encompass all genospecies of Lyme disease spirochetes, *B. burgdorferi* sensu lato.

**Figure 1.**
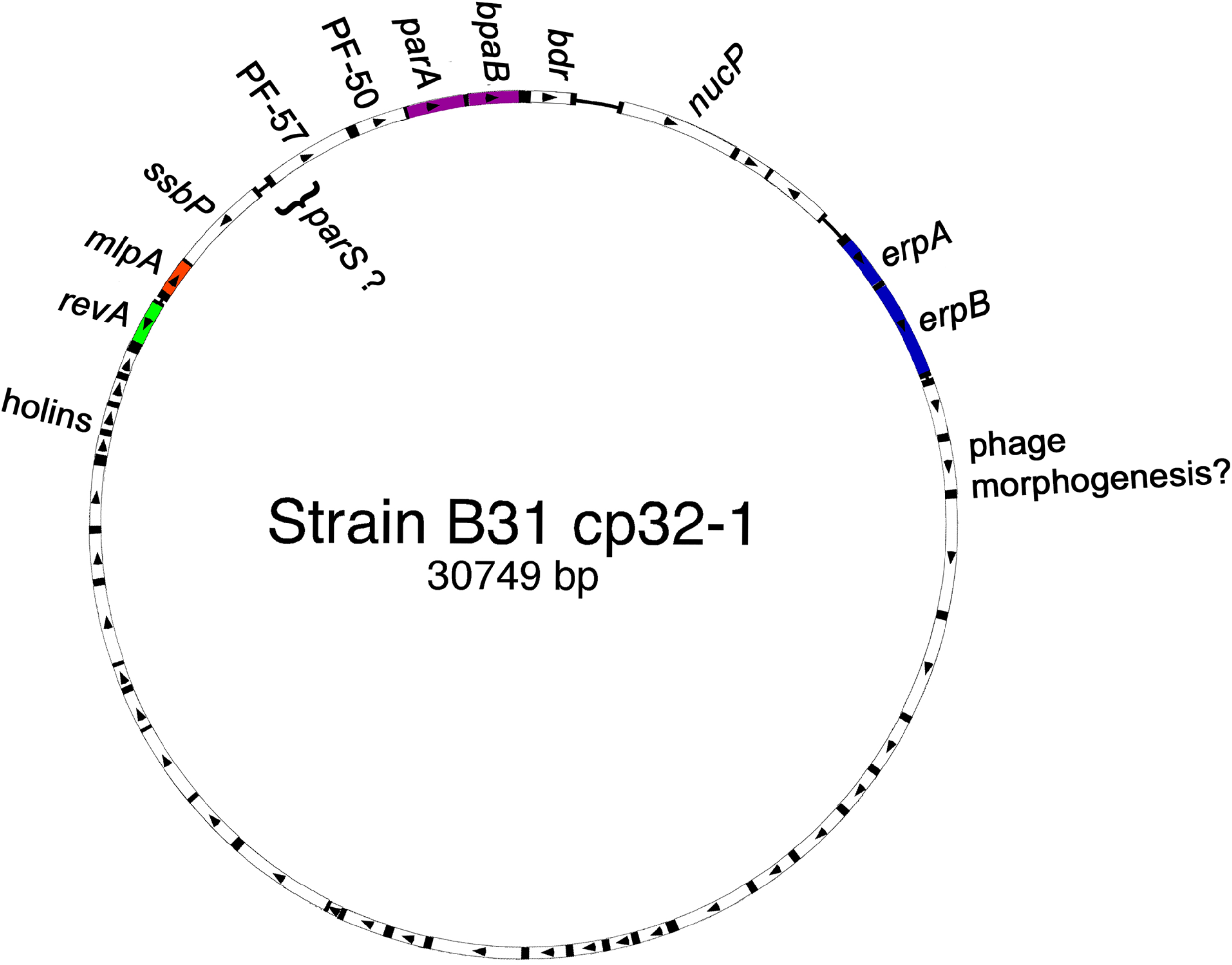
Schematic of the open reading frames of a representative *B. burgdorferi* cp32, the cp32-1 of type strain B31. Arrowheads indicate direction of open reading frame transcription. *B. burgdorferi* cp32s contain two loci that encode lipoproteins that are expressed on the bacterial outer surface: a mono-or bicistronic locus that encodes Erp proteins, and a separate locus that consists of either divergently-transcribed *revA* and *mlp* genes, or a *mlp* operon adjacent to a *bdr* gene. B31 cp32-1 and cp32-6 each carry a *revA* gene, while the other cp32s of that strain instead carry a *bdr* gene adjacent to their *mlp* locus. Borreliae are apparently able to maintain numerous, different cp32s in a single cell due to diversity among the *parA*-*bpaB* (<u>b</u>orrelial *<u>pa</u>r<u>B</u>*-analog) locus. Other open reading frames show up to 100% conservation between different cp32s and *B. burgdorferi* strains, shown as white. Two conserved genes, PF-50 and PF-57, are located adjacent to the *parA-bpaB* loci of all cp32s and are necessary for replicon maintenance (158-160). The *parS* segregation site appears to be associated with the 5’ end of *PF-57* (our unpublished results and (158-160). Conserved genes *ssbP* and *nucP* encode a single-stranded DNA binding protein and nuclease, respectively (35). Bacteriophage structural and assembly proteins appear to be encoded from a single operon that extends from 3’ of the *erp* locus to the holin-encoding genes (16, 39, 43, 161).

**Figure 2.**
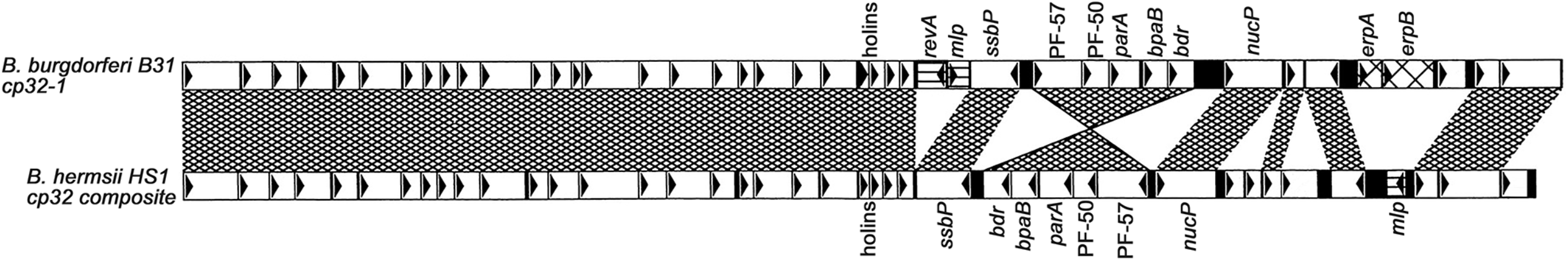
Alignment of the open reading frames of *B. burgdorferi* B31 cp32-1 and a composite cp32 of *B. hermsii* HS-1 that was assembled from sequenced contigs. Directions of transcription are indicated by arrows. The *erp, revA* and *mlp* genes are indicated by hatching. ORFs that are nearly identical between cp32s of *B. burgdorferi* and *B. hermsii* are indicated by white boxes. Note that the plasmid replication genes, including *parA* and *bpaB*, are located in reversed directions on the two different cp32 types. Adapted from (11).

*Borrelia* species all have multipartite genomes, consisting of an approximately one megabase main chromosome and numerous smaller replicons that are generally referred to as “plasmids” (14-16). Some borrelial “plasmids” carry essential genes, and might be better considered to be additional chromosomes (14, 16-20). Another uncommon feature of borrelial genomes is a preponderance of linear DNAs, with the main chromosome and many of the smaller replicons being linear elements with closed hairpin telomeres (14, 16, 18, 21-23). Lyme disease spirochetes and many other borreliae also maintain circular DNA replicons (24-26).

During the mid-1990s, evidence appeared that *B. burgdorferi* possesses multiple, homologous loci in (26-31). Further studies determined that each related locus is carried on a different circular replicon of approximately 32 kb in size, and that a single bacterial cell can contain numerous different variants of those replicons (9, 26). Based on the sizes and circular nature of the replicons, they were designated “cp32”s (<u>c</u>ircular <u>p</u>lasmid <u>32</u>kb). The majority of nucleotides that comprise different *B. burgdorferi* cp32s are highly conserved, differing significantly at only three loci: one that is responsible for plasmid maintenance and two that encode outer-surface lipoproteins (Fig. 1) (12). The highly repetitive nature of cp32s initially prevented assembly of their sequences, such that the initial, 1997 report of the genome of *B. burgdorferi* type strain B31 did not include the native cp32 plasmids (14). Genomics analysts subsequently used the cp32s as learning tools to refine sequence assembly algorithms, and the entire genome sequence of strain B31, including all cp32s and other repetitive sequences, was finally published in 2000 (16).

The different cp32s within a single bacterium appear to be compatible with each other due to their encoding two maintenance proteins, ParA and BpaB, that are distinct for each cp32 within a cell (31-36). Yet, *parA-bpaB* locus sequences are conserved between strains (13, 32, 36). Extensive characterization of cp32 sequences from Lyme disease spirochetes that were collected around the world revealed twelve apparent clades of cp32 *parA-bpaB* pairings (13, 36, 37). Additionally, some strains contain small plasmids of approximately 9 kb that were derived from cp32s and which carry only a *bpaB* allele (14, 16, 38). The absence of a *parA* gene from those cp9 plasmids raises intriguing questions about plasmid maintenance in *Borrelia* species.

The conserved size and sequence of *B. burgdorferi* cp32s, along with genes for proteins that resemble bacteriophage portals and holins, led to an early hypothesis that they might be prophage genomes (9, 39). Cultures of *B. burgdorferi* have been observed to clear spontaneously, or when exposed to 1-methyl-3-nitroso-nitroguanadine (MNNG), with production of bacteriophage particles that were named ϕBB1 (10, 40). Molecular analyses of those particles revealed that they contained cp32 DNA, and that they were capable of transducing nucleic acids between *B. burgdorferi* cells (41, 42). A subsequent study revealed that MNNG can induce transcription of a long, multigene cp32 operon that presumably encodes bacteriophage structural proteins (43). Intriguingly, even though MNNG is a mutagen, *B. burgdorferi* lacks an SOS system (14), so the means through which this chemical induces bacteriophage production is not yet known. It was recently reported that dysregulation of the RpoS alternative sigma factor led to production of ϕBB1 bacteriophage particles (44). Characterizing the mechanism that controls the lysogenic vs. lytic phases of ϕBB1 will undoubtedly yield important insights on the breadth of bacteriophage biology.

### *B. burgdorferi* cp32-encoded outer-surface Erp lipoproteins

Two antigenic outer-surface proteins of *B. burgdorferi* strain N40 were reported in 1994 and named OspE and OspF (outer surface proteins E and F) (45). Shortly afterward, we found that *B. burgdorferi* N40 expresses different levels of those two proteins under different culture conditions (46). Screening of a lambda library of *B. burgdorferi* type strain B31 DNA revealed several distinct loci in that strain that were closely related to, but distinct from, the strain N40 *ospE-ospF* locus (26). As none of the strain B31 loci could be definitively called “*osp<u>E</u>-ospF*”, they were designated “*erp*” (Osp<u>E</u>F-<u>r</u>elated <u>p</u>rotein). Further analyses revealed that various clonal cultures of *B. burgdorferi* B31 contain up to 9 different *erp* loci (9, 16) (Table 1). Strain N40 contains 6 *erp* loci, including the initially-described *ospE-ospF* (13, 36, 37, 45, 47, 48) (Table 1). Researchers who investigated other strains discovered additional *erp* genes, and gave them a variety of names such as *p21, bbk2*.*10, bbk2*.*11, pG* and *elp* (28, 29, 47, 49-53) (Table 1). Naming of these genes and proteins by multiple groups has, unfortunately, led to a rather chaotic nomenclature. Nonetheless, to avoid further confusion, we use the names originally applied to these genes and proteins, and urge other authors to also retain names that have been in common use for the past several decades. Structures have been determined for some Erp proteins, primarily alleles that occur in the B31 type strain (54-57).

**Table 1:**
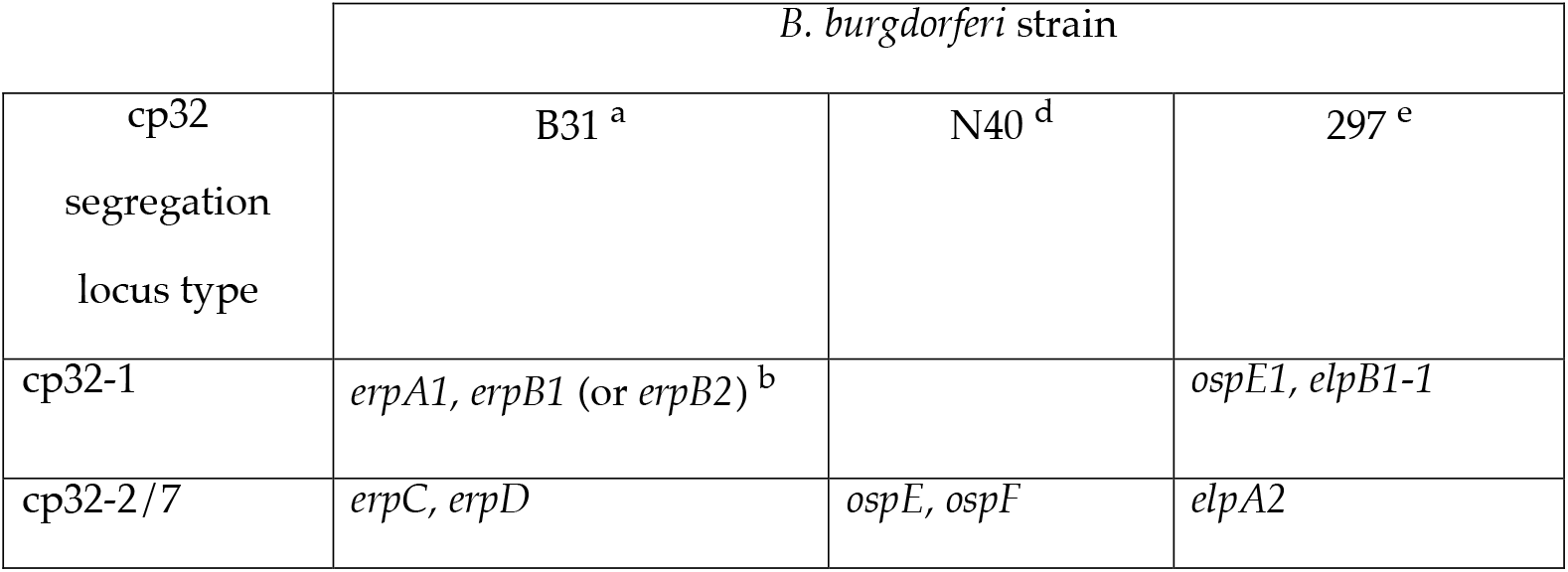
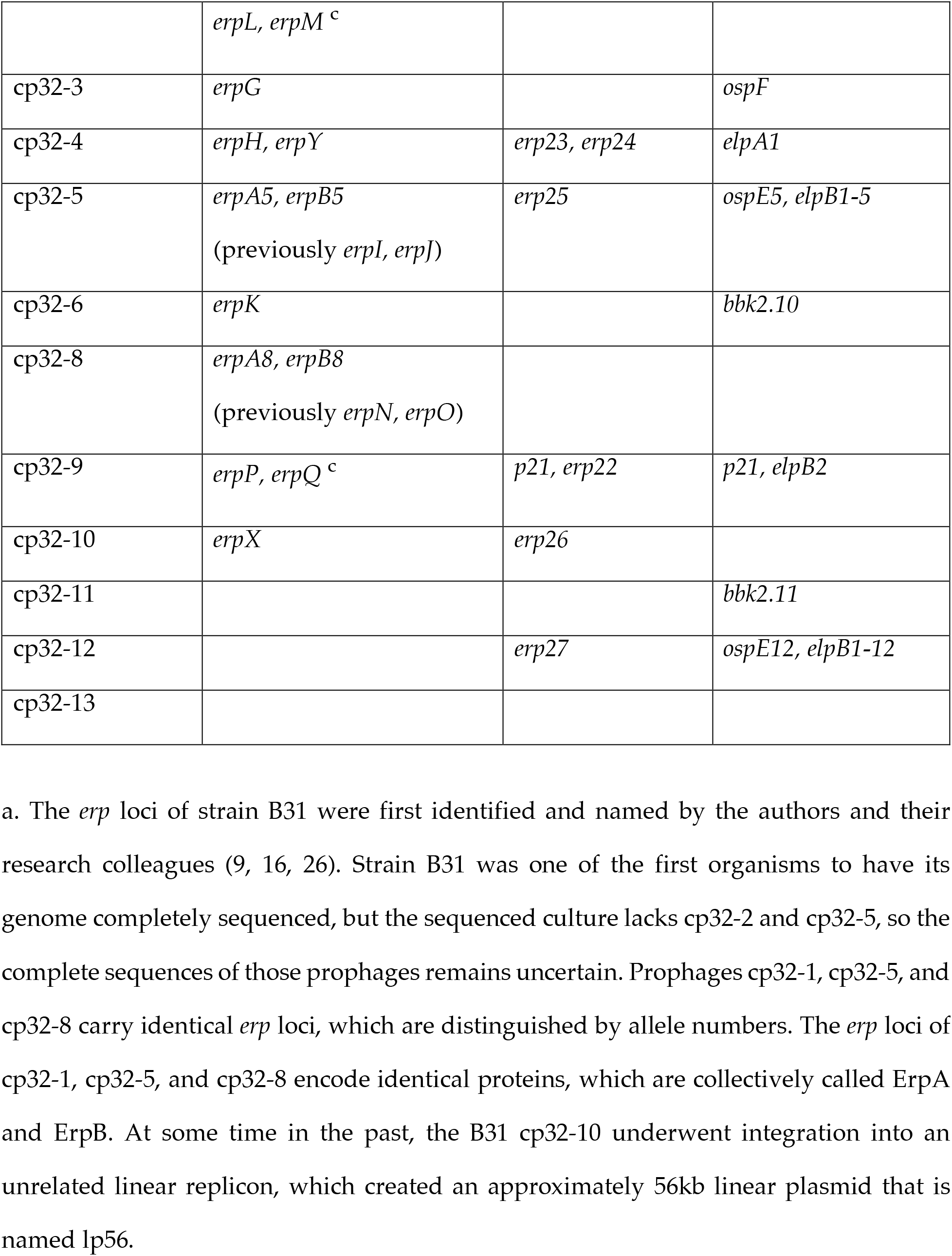
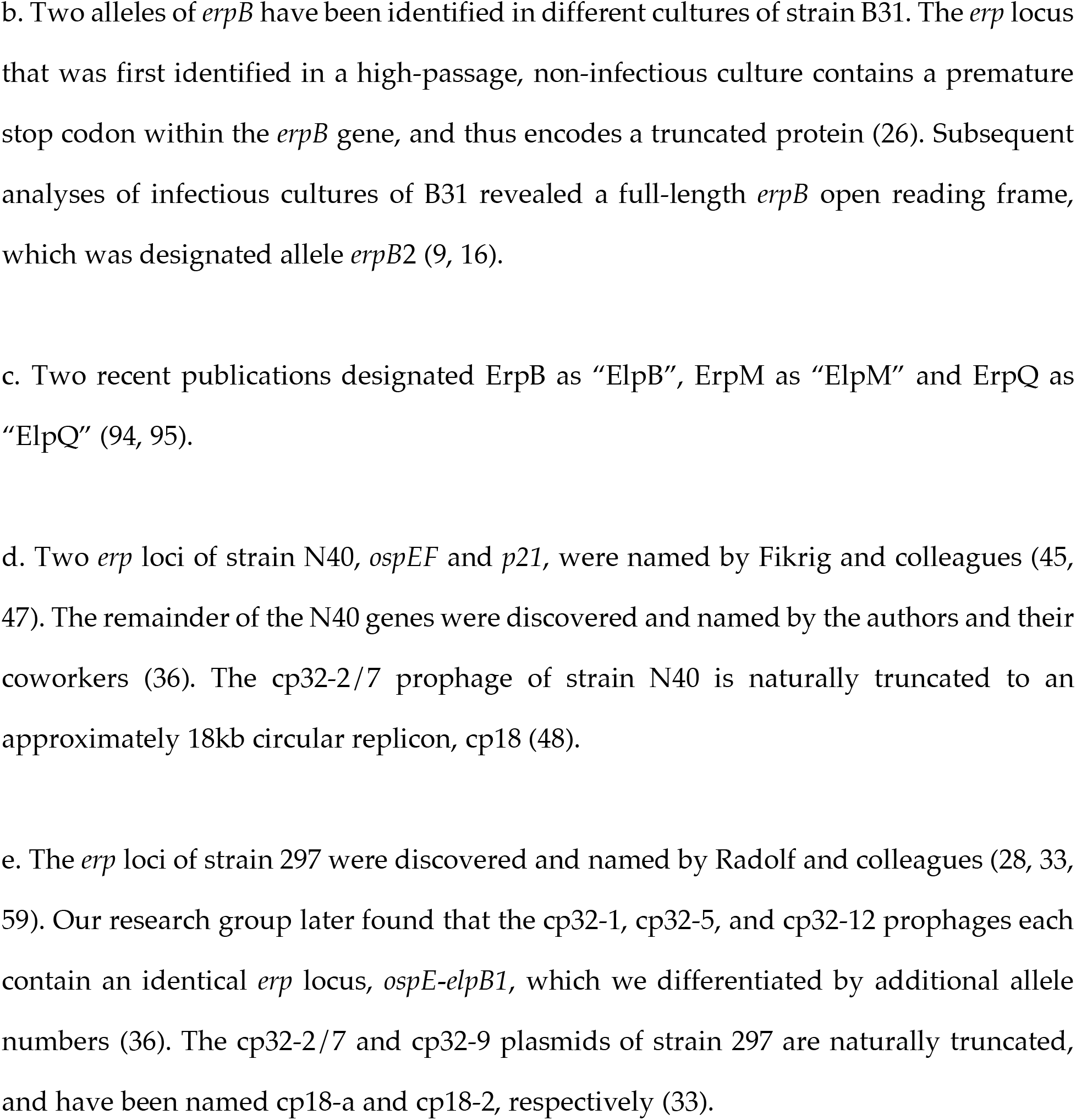
The cp32 DNA elements and associated *erp* loci that occur in the three best-characterized Lyme disease spirochete isolates: *B. burgdorferi* strains B31, N40, and 297. The *erp* nomenclature has been applied to the majority of other characterized strains of Lyme disease spirochetes (36, 88). Plasmid nomenclature is standardized according to segregation locus type, such that all plasmids designated “cp32-1” contain a similar segregation locus, etc. All known cp32 segregation loci fall into 12 clades. The first *B. burgdorferi* strain to be completely characterized, B31, contains 2 distinct cp32 prophages that possess identical maintenance loci, cp32-2 and cp32-7, and define segregation locus type 32-2/7. As a result, the numbering scheme for the 12 groups goes up to cp32-13. The published names of *erp* genes are listed. A variety of naming schemes were applied to these genes by their discoverers, particularly for strain 297, which has led to a sometimes confusing nomenclature. Additional *erp* genes of strain N40 that were discovered by our group were given allele numbers only (e.g. *erp22, erp23*, etc.). Other strain’s *erp* genes have been designated either by an allele number (with a different number for each gene) or by genome sequencing ORF numbers, which are too numerous to list here but examples are described in (36).

Considering *erp* genes and their proteins as a unified group is supported by their all occupying an allelic location on cp32s, their conserved promoter and operator DNA sequences, simultaneous co-expression in culture and during infection processes, possession of conserved leader polypeptides and initial residues of their mature lipoproteins, and localization of the mature lipoproteins on the outside of the borrelial outer membrane (12, 58). Evolutionary analyses indicated that *erp* genes arose from a single sequence that encodes the leader polypeptide and the 5 amino acid sorting sequence, while the remainder of the open reading frames descended from a large number of independent origins that frequently undergo novel recombination events (36). Dendrograms of known *erp* sequences show at least three branches (32, 36, 59), which led to a suggestion that *erp* genes could be divided and given three different group names: *ospE, ospF*, and *elp* (59). In addition to variations in nucleotide sequences, the encoded proteins generally differ considerably in size, with OspE-types being small (approximately 20 kDa), OspF-types being medium sized (approximately 25-30 kDa), and Elp-types being large (approximately 40 kDa) (36). We do not find the OspE/OspF/Elp nomenclature to be particularly helpful, since functions are not always conserved within each group, and there are no consistent differences in gene expression or operon structures. In addition, due to frequent natural recombination events having occurred between *erp* genes, some extended nucleotide sequences are found in members of all three groups (36). For example, some “*elp*” alleles share greater similarities with “*ospF*” genes (as high as 34%) than they do with other “*elp*” genes (as low as 17%) (36). Recombination events have also resulted in alleles that do not fit in the tripartite scheme (36). Extensive analyses of *erp* open reading frames, operon structures, promoter sequences, and both intragenic and intergenic recombination, indicated that *erp* genes evolved from a single promoter/leader/localization sequence and a large number of diverse sequences that encode the external parts of the proteins (36).

Erp lipoproteins are localized to the borrelial outer surface (45, 60, 61). The strain B31 Erp proteins are all simultaneously co-expressed, consistent with their genes possessing conserved promoter and operator sequences (62-64). Differences in gene or protein expression have been reported for two other strains, but the reasons for those apparent variations were not investigated (47, 61, 65). The mechanism by which *B. burgdorferi* controls transcription of *erp* operons has been well studied, and shown to involve interactions between the BpaB repressor, BpuR co-repressor, and EbfC antirepressor proteins that bind to specific sites adjacent to *erp* promoters. There is evidence that some *erp* operons may be affected by an alternative sigma factor, RpoS (66, 67). For an extensive review of the mechanisms by which *erp* transcription is regulated, please see our recent review on that topic (68).

*B. burgdorferi* expresses little to none of any Erp protein during colonization of unfed ticks (47, 63, 69). As the tick begins to feed on the blood of a vertebrate host, the bacteria induce production of their entire repertoire of Erp proteins (63). Indirect fluorescence analyses (IFAs) of skin samples at the sites of tick attachment revealed that almost all transmitted *B. burgdorferi* produced detectable levels of every examined Erp protein (63). Quantitative reverse transcription PCR (qRT-PCR) and promoter fusions with *gfp* revealed that *erp* genes remain induced throughout persistent mammalian infection (53, 63, 70). Levels of antibodies against Erp proteins, including IgM, remain elevated during prolonged infection of mice (53, 63, 70-72). The persistence of IgM antibodies against Erp proteins may reflect *B. burgdorferi*’s ability to manipulate host immune responses (73-76). That phenomenon may also explain how *B. burgdorferi* is able to persist extracellularly despite having its outer surface covered with Erp, Rev, and other antigenic proteins. IFAs of naive ticks feeding on infected mice showed that *B. burgdorferi* continue to produce Erp proteins during acquisition by feeding ticks, which ceases soon after detachment (63).

Numerous functions have been identified for Erp proteins, all of which involve interactions with vertebrate host tissues:

Several Erp proteins, including ErpA, ErpC, and ErpP of strain B31 and OspE of strain N40, exhibit high affinities for complement factor H and the related proteins FHL-1 and the FHR group (71, 77-91). Binding factor H and FHL-1 to *B. burgdorferi* via Erp proteins confers resistance to the alternative pathway of complement activation. The exact functions of FHR proteins are unclear, but they also appear to be involved with complement control (92). Erp proteins with different sequences show varying affinities to factor H molecules of different potential vertebrate host species, and may thus play roles in expanding the host range of Lyme disease spirochetes (85, 93).

The ErpB, ErpM and ErpQ proteins of strain B31 (confusingly called ElpB, ElpM and ElpQ in the publications) were recently demonstrated to bind complement factors C1s and C1r (94, 95). Adhesion of those host proteins onto the borrelial surface appears to afford protection from the classical arm of complement.

The ErpX protein of *B. burgdorferi* B31 binds human laminin (96). This may help the spirochete target and/or adhere to laminin-rich tissues. The laminin-binding domain of ErpX was mapped to a central, disordered domain that is almost entirely comprised of charged amino acids (96). Of note, other strain B31 Erp proteins with amino acid sequences that are similar to ErpX (others in the “Elp” group: ErpB, ErpM and ErpQ, Table 2) do not detectably bind human laminin (96).

**Table 2.** The Erp proteins of the three best-characterized *B. burgdorferi* strains, B31, N40, and 297, sorted according to the tripartite OspE/OspF/Elp scheme. Other characterized strains of Lyme disease spirochetes have generally applied the *erp* nomenclature (36, 88). Please note that even though some proteins of strain 297 proteins were assigned the same names as N40 proteins, those proteins are distinct from each other. That is, N40 OspE is not identical to 297 OspE, N40 P21 is not identical to 297 P21, and N40 OspF is not identical to 297 OspF. The strain 297 nomenclature is further complicated in that the 297 OspE protein is more similar to the N40 P21 protein than to the N40 OspE protein, and the 297 P21 protein is more similar to the N40 OspE protein (36).

|  | “OspE”<br>“small”<br>(69% average<br>pairwise identity) <sup>a</sup> | “OspF”<br>“medium”<br>(44% average<br>pairwise identity) <sup>a</sup> | “Elp”<br>“large”<br>(40% average<br>pairwise identity) <sup>a</sup> |
| --- | --- | --- | --- |
| *B. burgdorferi* B31 | ErpA<br>ErpC<br>ErpH<br>ErpP | ErpG<br>ErpL<br>ErpK<br>ErpY | ErpB<br>ErpD<br>ErpM<br>ErpQ<br>ErpX |
| *B. burgdorferi* N40 | OspE<br>P21 | OspF<br>Erp23<br>Erp25<br>Erp27 | Erp22<br>Erp24<br>Erp26 |
| *B. burgdorferi* 297 | OspE<br>P21 | OspF<br>Bbk2.10 | ElpA1<br>ElpA2 |
|  |  | Bbk2.11 | ElpB1<br>ElpB2 |

a. Average pairwise identities of genes and proteins of 21 analyzed Lyme disease *Borrelia* strains (36). That study found that some alleles that fall into the OspF group due to their sizes share greater sequence identity with members of the Elp group than they do with other members of the OspF group, and vice versa. Those similarities and differences appear to be due to frequent recombination between alleles of different groups.

The *B. burgdorferi* B31 ErpG and six similar proteins from other strains bind heparan sulfate, and promote attachment to glial cells (97). A phage display library, injected into mice and screened for potential *B. burgdorferi* adhesins, found clones of B31 ErpG, ErpK, and ErpL, and strain 297 Bbk2.10 in mouse heart, bladder, and tibiotarsal joint tissues (98). Those four proteins share some sequence similarities (Table 2) (36).

The *B. burgdorferi* B31 ErpA, ErpC, and ErpP proteins bind host plasminogen and its proteolytically-active form, plasmin (99). Plasmin bound to Erps and other surface proteins facilitates *B. burgdorferi* migration through solid tissues in vitro, a characteristic that is hypothesized to also enable dissemination during vertebrate infection (100-109). It is notable that the Erp proteins that bind plasmin(ogen) also bind complement factor H (see above).

Due to the surface localization and antigenicity of Erp proteins, there have been several investigations into their use as protective vaccines or for serological diagnosis of *B. burgdorferi* infection. However, vaccination of mice with the OspE, OspF, or P21 proteins of strain N40 did not provide substantial protection against infection by the same strain (47, 69). Several recombinant Erp proteins have been examined for serological diagnosis of human infection, with mixed results (110-113).

### *B. burgdorferi* cp32-encoded outer-surface Rev and Mlp lipoproteins

The cp32 replicons of Lyme disease and relapsing fever borreliae carry a distinct mono-or bicistronic *mlp* (multicopy lipoprotein) operon (Figs. 1 and 2) (11, 16, 30). In the majority of Lyme disease spirochete cp32s, the *mlp* locus is preceded by a distinct gene, *bdr*, which has its own promoter (16, 30) (Fig. 1). The characterized cp32 of the relapsing fever spirochete *B. hermsii* contains a single *mlp* gene, without an adjacent *bdr* (11). Curiously, all cp32s of Lyme disease and relapsing fever borrelia carry a second *bdr* allele adjacent to the episomal replication locus (9, 16, 31) (Fig. 1). Bdr proteins are associated with the bacterial inner membrane, are produced during vertebrate infection, but their function(s) have yet to be discovered (72, 114-122).

For unknown reasons, some *B. burgdorferi* cp32s do not contain a *bdr* adjacent to the *mlp* locus, but instead carry a distinct gene named *revA* (so named because such genes are arranged reverse to *mlp*) (16, 30). The *revA* ORF and promoter sequences are very different from those of *mlp* genes, implying that *mlp* and *revA* loci are genetically unrelated (9, 16, 30). In the *B. burgdorferi* type strain, B31, cp32-1 and cp32-6 each contain a *revA* gene adjacent to their *mlp*, while all other B31 cp32s contain only a *mlp* gene (16).

Mature RevA is a lipoprotein and localizes to the *B. burgdorferi* outer surface (123). Additionally, some truncated cp32 derivatives, such as B31 cp9, carry a similar gene named *revB* (14, 16, 123). The two *revA* orthologs of strain B31 encode an identical RevA protein, while that strain’s RevB protein shares only 28% amino acid identity with RevA. Other strains encode RevA and RevB proteins with varying degrees of differences (30).

*B. burgdorferi* in unfed ticks do not produce detectable levels of *revA* mRNA, but do produce elevated transcripts during mouse infection (124). Human patients and experimentally infected mice produce antibodies against RevA (125-127). In culture, *revA* transcription is induced by changes in temperature or pH that correspond to changing conditions encountered by the bacteria during tick feeding and mammalian infection (123). Despite its antigenicity and surface localization, vaccination with RevA did not protect mice from *B. burgdorferi* infection (128).

*B. burgdorferi* B31 RevA binds the 70kDa N-terminal region of fibronectin via a domain in the RevA N-terminus (124). This binding was not affected by salt or heparin, suggesting the interaction of these two proteins involves more than ionic interactions. The *B. burgdorferi* RevB protein also binds to fibronectin, despite having limited sequence identity to RevA (124). Polyclonal antibodies against RevA did not block adherence of fibronectin to wild type *B. burgdorferi*, although those antibodies did prevent fibronectin adherence to *B. burgdorferi* that were deleted of another fibronectin-binding protein, BBK32 (129, 130). Similarly, binding of wild-type *B. burgdorferi* to HUVECs or human neuroglial cells was not blocked by anti-RevA monoclonal antibodies (131). Those results underscore the functional redundancy of *B. burgdorferi* fibronectin-binding proteins (132). RevA also binds mammalian laminin, although to a significantly lesser extent than to fibronectin (124).

A *B. burgdorferi* mutant unable to produce RevA was found to be defective in colonization of cardiac tissue (130). That mutant also caused increased pathology of tibiotarsal joints and enhanced CCL2 production. Increased inflammation in the absence of RevA protein suggests a role for that surface protein in immune evasion, perhaps by masking a more immunogenic antigen.

Precise functions have not yet been ascribed to Mlp proteins. The cp32 prophages of both Lyme disease and relapsing fever *Borrelia* spp. carry *mlp* genes, which suggests that the proteins serve functions that are conserved among borreliae (9, 11, 16, 30, 133) (Fig. 2). As with *erp* and *rev* genes, there is often considerable sequence diversity among *mlp* genes within and between *B. burgdorferi* isolates (16, 133, 134). The patterns of regulated expression and antigenicity of *B. burgdorferi* Mlp proteins imply that they serve functions within vertebrate hosts (134-139). A recombinant Mlp protein was found to adhere to human brain microvascular endothelial cells (140). The structure of the MlpA/P28 protein that is encoded by B31 cp32-1 has been solved, and was noted to contain a structural motif that is similar to one seen in a lipid-binding protein (16, 134, 141).

### Horizontal transfer and recombination

Almost all examined Lyme disease spirochete isolates naturally contain 6-12 distinct cp32s, each with an *erp* and a *mlp* / *revA* locus. As noted above, relationships between cp32s can be discriminated by their *parA-bpaB* maintenance loci. Using that as reference, there is clear evidence of recombination having occurred between cp32s, and exchange of cp32 DNA between bacteria (13, 32, 36, 37).

The original isolate of *B. burgdorferi* type strain B31 carried three identical *erpAB* operons on cp32-1, cp32-5, and cp32-8 (9, 26). In contrast, those cp32s of other strains possess distinct *erp* operons (32, 33, 36). The cp32-1 and cp32-6 of strain B31 carry *revA* genes that are nearly identical in sequence, and which encode identical RevA proteins, while related plasmids of other strains have different genes at that locus (16, 124, 127). Two other recombination events are also evident in strain B31: cp32-4 has an fragment of a cp32 *sbbP* gene inserted into *erpH*, such that a complete lipoprotein cannot be produced, and cp32-3 has an unrelated gene, *bapA*, inserted 3’ of *erpG* (9, 26). No other B31 cp32 carries a *bapA* gene, although paralogous genes, named *eppA*, are found on some cp9 replicons (142-144).

Despite cp32s having nearly identical sequences, and multiple cp32s residing within individual bacteria, there are no known examples of recombination occurring during cultivation. Recombination does not take place during vertebrate infection, or, at least, not among bacteria of the same strain (145, 146). A report suggesting that variation arose during mammalian infection was subsequently seen to be based on PCR artifacts (146, 147). We hypothesize that genetic transfer and recombination occur when *Borrelia* spirochetes are within ticks. A recent study of borrelia bacteriophage induction supports that hypothesis (44).

Additional evidence of horizontal transfer of cp32 DNAs in nature was obtained from sequencing genomes of multiple isolates (36). *B. burgdorferi* strains B31 and BL206 contain almost identical repertoires of cp32s, the only differences being presence of a cp32-1 in only B31 and a cp32-11 in only BL206. Intriguingly, B31 cp32-1 and BL206 cp32-11 carry an identical *erpAB* locus. Two other strains, 297 and Sh-2-82, contain cp32 repertoires that are identical to each other, with the addition of a cp32-8 in only Sh-2-82. The cp32-8 *erp* locus of Sh-2-82 is identical to the cp32-8 *erpAB* locus of strain B31, implying that a cp32-8/ϕBB1 was exchanged between ancestors of those two strains. Strain B31 was isolated from a tick on Shelter Island, New York, in 1981, and Sh-2-82 was isolated from another tick on the same island in 1982 (148, 149). It is also notable that BL206, which is a near clone of B31, was isolated several years later from the blood of a Lyme disease patient in Westchester County, New York (150), and that 297, which is a near clone of Sh-2-82, was isolated from the cerebrospinal fluid of another patient (151).

Transduction of cp32 DNA between different strains of *B. burgdorferi* has been observed in culture (41). The mechanisms that induce ϕBB1 particles to form are not yet known. As noted above, *B. burgdorferi* does not possess an SOS system (14). The timing of bacteriophage production is also unknown. All *Borrelia* species are obligate parasites of arthropods and vertebrates, so triggers for ϕBB1 would be limited to conditions experienced within those hosts. Ticks can acquire multiple strains of *B. burgdorferi*, which would reside together in the tick’s midgut, leading us to propose that the midguts of feeding ticks are likely locations for DNA exchange. Tick feeding is a time of rapid replication for *B. burgdorferi*, which appears to be used by the spirochetes as an important cue that transmission is imminent (68, 152). That would also be an appropriate time for bacteria to exchange DNA and diversity their repertoires of Erp, Rev, and other surface proteins that interact with vertebrate hosts.

At least two advantages appear to be conferred upon *B. burgdorferi* by exchanging and recombining *erp* and *rev* genes. First, both groups of proteins are expressed early during vertebrate infection, are located on the bacterial outer surface, and are antigenic. Reservoir vertebrates are generally fed upon by numerous ticks, all of which might be infected. Thus, Lyme disease spirochetes are under selective pressure to diversify sequences of early, surface-borne antigens, to facilitate infection of animals that have previously been exposed. A second advantage to Erp and Rev variability among individual bacteria stems from their functions as adhesins, and from the fact that the ticks that transmit Lyme borreliae may not be specific in their choices of hosts. For example, in the northeastern USA, *Ixodes scapularis* ticks feed on birds, mice, and other mammals (153). Since a *B. burgdorferi* cannot “know” in advance what species of vertebrate its tick vector will feed upon, it would be advantageous for a bacterium to enter its new host while bristling with a wide variety of adhesins that differ in relative affinities for ligands of different hosts, with the likelihood that some Erp, Rev, and other adhesins will be adequate for the bacteria’s survival.

### Exceptional *Borrelia*

To date, only two exceptions are known to the generalization that *Borrelia* species naturally carry cp32s. The agent of avian spirochetosis, *Borrelia anserina*, is the type species of the genus, and lacks cp32 DNA (11). Since Lyme disease and relapsing fever *Borrelia* species carry cp32s, their absence from *B. anserina* suggests that either ϕBB1 entered in the genus after the split of *B. anserina* from the other species, *B. anserina* has a means to exclude cp32s/ϕBB1, or cp32s encode properties that are beneficial to Lyme disease and relapsing fever borreliae, but not to avian spirochetosis borreliae. The second known example of a cp32-deficient *Borrelia* is strain Far04, a Lyme disease spirochete that was isolated from a puffin that was collected on the Faroe Islands, Denmark (13, 154, 155). The densely packed puffin nests on those islands are homes to *Ixodes uriae* ticks, which preferentially feed on seabirds. Although Far04 is a single example, it is possible that *I. uriae* ticks on the islands primarily feed on puffins, to the extent that some circulating borreliae have lost the need for the variability of adhesin compositions afforded by cp32 prophages. Clearly, additional sampling and analyses are needed to test that hypothesis.

### Future research directions

Erp and Rev lipoproteins localize to the borrelial outer surface, and are known to interact with a variety of vertebrate host proteins that protect against immune responses and facilitate tissue invasion and colonization. Noting that Erp and Rev proteins were discovered during the mid-1990s, and that adherence of complement C1r and C1s was first reported in 2022, we consider it likely that other functions remain to be discovered. Additional questions include: Why do the cp32s of Lyme disease spirochetes encode Erp and Rev proteins, while those of relapsing fever spirochetes appear to lack those genes? What are the functions of the Mlp outer surface proteins, which are encoded on cp32s of both Lyme disease and relapsing fever borreliae? Why does *B. anserina*, the agent of avian spirochetosis, lack cp32s? When do borreliae produce ϕBB1 bacteriophages during their natural infectious cycles? There is genetic evidence of DNA transfer of other genes encoding borrelial antigens, such as outer surface protein C (OspC) (156, 157) – might that occur by generalized transduction via ϕBB1?

## Concluding statements

The cp32 prophages / ϕBB1 bacteriophages have evidently co-evolved with many species of *Borrelia*. They have acquired a number of different genes that confer benefits to their bacterial hosts, which thereby enhance their own survival. These bacteriophages have adopted a primarily lysogenic lifestyle, with a limited degree of bacteriophage particle production and host cell lysis. The *parA* and *bpaB* maintenance genes have diversified extensively, to the extent that large numbers of different cp32s can be maintained in a single bacterial cell. This provides both extensive repertoires of *erp, rev*, and other genes for horizontal exchange and enables the host bacteria to express wide varieties of surface adhesins during vertebrate infection.

## Acknowledgements

We thank the many colleagues from our laboratories and throughout the world who have contributed to the studies described in this review. Research in our laboratories is currently funded by NIH grants R01AI144126, 3R01AI144126-03S1, and R21AI147139 to Brian Stevenson and R01AI58304 to Catherine Brissette.

